# Natural variation in phospholipase IPLA2-VIA contributes to starvation sensitivity in *Drosophila melanogaster*

**DOI:** 10.1101/2024.07.05.602254

**Authors:** Shawn B. Williams, Brian Franklin, Faye A. Lemieux, David M Rand

**Author notes:** Correspondence: David M Rand, Brown University, Box G-W, 80 Waterman Street, Providence, RI 02912. Current address: University of Pennsylvania Perelman School of Medicine, Philadelphia, PA 19104, USA.

## Abstract

Resistance to starvation is a classic complex trait where genetic and environmental variables can greatly modify an animal’s ability to survive without nutrients. Genetic analyses in *Drosophila* have shown that starvation resistance is highly polygenic with different genetic architectures in different mapping populations. In this study we sought to dissect the genetic basis of starvation resistance among a set of mitonuclear genotypes carrying different mtDNAs paired with specific nuclear genomes from the Drosophila Genetic Reference Panel (DGRP). We focused on differences between one of the most sensitive strains (DGRP-765) and a strain with more moderate resistance (DGRP-315) whose starvation phenotypes appeared to be modified by alternative mtDNAs. Using complementary pooled-sequencing and forward genetic mapping approaches, we identified regions of chromosomes 2L, 3L and 3R contributing to starvation sensitivity and localize a major effect locus modifying starvation resistance to the coding region of phospholipase iPLA2-VIA. These analyses further confirm that the alternative mtDNAs had little influence on variation in starvation resistance between the genotypes studied. The sensitive line shows a starvation-dependent depletion of glucose and glycogen that is modified by hemi- and heterozygosity in the iPLA2-VIA region. These findings contribute to our understanding of the complex genetic relationship between resistance to starvation stress and nutrient metabolism.

## Introduction

Coping with periods of low nutrient availability is a fundamental fitness component of all organisms. In animals, multiple metabolic processes are required to convert dietary nutrients into usable resources to successfully cope with periods of starvation. Energy that is not needed immediately has to be converted into a form amenable for efficient storage and that stored energy has to be accessible for utilization during periods of starvation. For these reasons, the genetics of starvation resistance provides a compelling model for understanding the integrative aspects of animal physiology (Brown *et al*. 2019). Furthermore, a better understanding of the genetic basis of nutrient catabolism and storage will also shed a light on genes relevant to metabolic diseases and fitness traits in nature (Hardy *et al*. 2018).

Much of the information on the genetics of starvation resistance comes from studies in Drosophila employing either direct selection on experimental outbred populations or genome wide association studies (GWAS) across samples of inbred lines. Variation in starvation resistance is an appealing trait for studies of quantitative and evolutionary genetics due to its highly polygenic nature and its correlation with other fitness and metabolic traits (Service and Rose 1985; Chippindale *et al*. 1996; Harshman *et al*. 1999; Hardy *et al*. 2018; Everman *et al*. 2019). GWAS approaches using either the Drosophila Genetic Reference Panel (DGRP;(Mackay *et al*. 2012)) or the Drosophila Synthetic Population Resource (DSPR;(Everman *et al*. 2019)) have identified a number of candidate loci influencing variation in starvation resistance confirming the complexity of its polygenic nature (Harbison *et al*. 2004; Sorensen *et al*. 2007; Huang *et al*. 2012; Mackay *et al*. 2012; Everman and Morgan 2018; Everman *et al*. 2019). Importantly, there are clear differences among studies in the location of QTLs and the putative candidate loci identified (Huang *et al*. 2012; Mackay *et al*. 2012; Everman and Morgan 2018; Everman *et al*. 2019). These differences have been attributed to the distinct genetic composition of the mapping populations and presumably unique epistatic interactions among loci influencing starvation resistance (*ops. cit.*). Notably, none of the QTL identified in the two subsets of the mapping lines used by (Everman *et al*. 2019) overlapped, likely due to the different founder lines used to build each subset (DSPR pA and pB). Despite these differences, there are shared loci that emerge from a comparison of multiple studies (table S9 in (Everman *et al*. 2019)).

A growing literature has identified a role for mitochondrial functions in stress responses at multiple levels (Uma Naresh *et al*. 2022; Erkosar *et al*. 2025). Given the central roles that mitochondria play in both anabolic and catabolic functions, the complex interactions between mitochondrial and nuclear genomes may play important roles in adaptive traits, and particularly in metabolically demanding conditions such as starvation, oxidative stress and animal performance (Rand *et al*. 2004; Hill 2015). In previous work, we demonstrated the impact of mitonuclear epistasis in development time among a panel of 72 mitonuclear genotypes constructed from 6 mtDNAs introgressed into 12 nuclear genomes from the Drosophila Genetic Reference Panel (DGRP) (Mossman *et al*. 2016). We selected a subset of these lines to examine starvation resistance based on significant differences in development time between mtDNAs in each of several DGRP backgrounds. These analyses revealed a putative mitonuclear epistatic interaction influencing starvation resistance and its relation to diet composition. In the current study, we sought to use a combination of quantitative genetic and classical genetic mapping approaches to examine the genetic basis of starvation resistance between two DGRP lines carrying different mtDNAs (DGRP-765 and -315). Genomic sequencing of sensitive and resistant pools of adults from advanced intercross populations derived from these lines identified regions of allelic differentiation. Chromosome segregation and recombination mapping identified major factors on chromosome 3L, 3R and 2L, but not chromosome X, 2R or the mitochondrial genome. Deficiency mapping indicated that the starvation sensitivity in DGRP-315 localizes in part to the phospholipase iPLA2-VIA gene with associated effects of carbohydrate metabolism. The correspondence of these approaches contributes to our understanding of the polygenic nature of starvation resistance and points to polymorphisms in the iPLA2-VIA region that motivate further study of the associations between starvation sensitivity, lipid metabolism and carbohydrate metabolism.

## Materials and Methods

### Fly stocks

The current study focuses on the differences in starvation resistance between specific mitonuclear genotypes carrying alternative mtDNAs placed onto nuclear genomic backgrounds of the Drosophila Genetic Reference Panel (DGRP;(Mackay *et al*. 2012)). In a previous study we constructed a panel of 72 mitonuclear genotypes using balancer chromosome substitution to place six mtDNAs onto each of 12 DGRP nuclear backgrounds to study mitonuclear epistasis for development time and its dietary modification (Mossman *et al*. 2016). From these 72 lines we quantified starvation resistance of 12 mitonuclear genotypes consisting of four different mtDNAs placed on three DGRP nuclear backgrounds (lines DGRP-765, -315 and -820). The four mtDNAs included two from *D. simulans* (*siI and siII*) and two from *D. melanogaster* (*OregonR* = *OreR* and *Zimbabwe53 = Zim53*). These mtDNAs were chosen to test for the impact of mtDNA polymorphism and divergence on starvation resistance. The DGRP backgrounds were chosen based on published starvation data showing that DGRP-765, -315 and -820 had low, medium and high values of starvation resistance, respectively (Ayroles *et al*. 2009; Mackay *et al*. 2012). The genotype notation used is *mtDNA;nuclearDNA*, e.g., *Zim53;315* carries the *D. melanogaster Zim53* mtDNA on a homozygous *DGRP-3*15 chromosomal background.

The results of this starvation assay are shown in Figure 1. The resistance levels were consistent with previously published rankings of the three nuclear genotypes (820 > 315 > 765). The results further showed that one unique combination, *siI;765,* (a *D. simulans siI* mtDNA on the *DGRP-765* nuclear background) had a significant increase in starvation resistance compared to the other mitonuclear genotypes in the *DGRP-765* background. The goal of the current study was to determine whether the apparent ‘rescue’ of starvation resistance was due to the introgression of the *siI* mitochondrial haplotype or due to a nuclear factor or factors. The focal mitonuclear genotypes are shown in Table 1. Additional genotypes for segregation, recombination and deficiency mapping are listed in Table S1.

**Figure 1.**
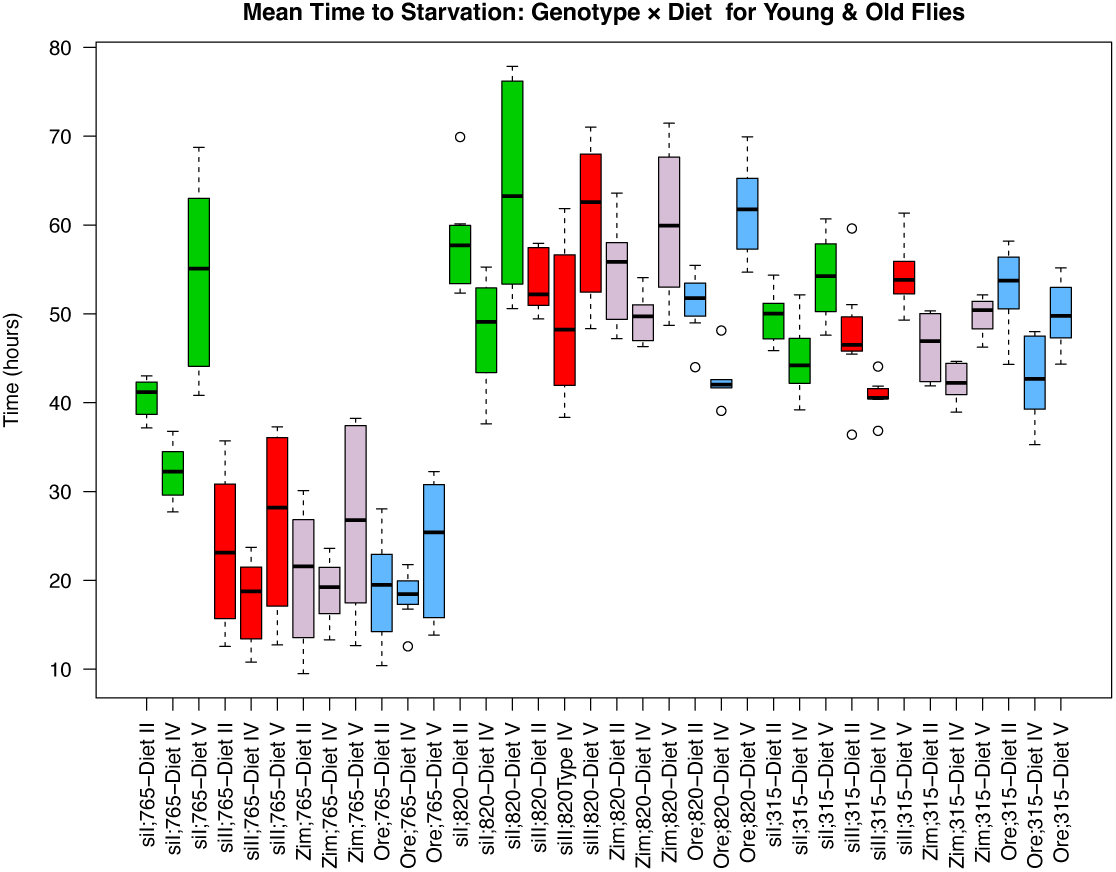
Starvation time among mitonuclear genotypes on different diets. The x-axis lists the genotypes with the notation: mtDNA;nucDNA-Diet, with Diets I, IV and V being balanced, low and high sugar:yeast ratios, respectively (10g:10g, 5g:15g, 15g:5g; see Methods).

**Table 1.** Mitonuclear stocks used for genetic analyses in this study. A table of additional stocks used for chromosome substitution, recombination mapping and deficiency mapping are listed in supplemental file 1.

| Stock Name<br>(mtDNA;nuclearDNA) | Mitochondrial<br>haplotype | Nuclear haplotype | Phenotype |
| --- | --- | --- | --- |
| *Zim53;765* | *D. melanogaster*<br>*Zim53* | DGRP-765 | Starvation<br>sensitive |
| *sil;765* | *D. simulans sil* | DGRP-765<br>recombinant | Starvation<br>resistant |
| DGRP-765 | From original<br>DGRP-765 line | DGRP-765 | Starvation<br>sensitive |
| *Zim53;315* | *D. melanogaster*<br>*Zim53* | DGRP-315 | Starvation<br>resistant |

### Starvation Stress

Survival during starvation stress was measured by placing approximately 125 female flies per genotype in each of three replicate demography cages. Flies were collected and sorted over CO_2_ and allowed to recover in quart-sized plastic demography cages affixed with a 25×95 mm vial containing one of three different isocaloric food media: high sugar food (15% sucrose, 5% Yeast, 2% agar), balanced food (10% sucrose, 10% yeast, 2% agar) or low sugar food (5% sucrose, 15% yeast, 2% agar; referred to as food type V, II and IV, respectively, in (Zhu *et al*. 2014)). The three replicate cages for each genotype were conditioned on these diets for 7 days unless otherwise indicated, with new food vials added every other day. After 7 days, any dead flies were removed, and the vial for each demography cage was replaced with 2% Difco Bacto-agar prepared with deionized H_2_0 to initiation starvation. The number of dead flies was recorded every 12 hours and removed from each demography cage until all flies were dead. As starvation resistance values for the different genotypes were most distinct on the high sugar diet “V”, most of the genetic mapping was carried out on that diet. The survival data from each cage were generally consistent so counts from each replicate cage were pooled and the combined data modeled using a Kaplan-Meir survival model implemented in R (Therneau and Grambsch 2000). Significant differences in survivals were determined using a log rank test.

### Forward genetic mapping of starvation sensitivity

*Chromosomal localization of resistance*. To determine if different mtDNAs were associated with the difference in starvation resistance reciprocal crosses were done between siI;765 and Zim53;765 as follows: virgin female siI;765 x male Zim53;765 and virgin female Zim53;765 x male siI;765). Maternal inheritance of mtDNA provides a test for the role of the *siI* mtDNA in the rescue of starvation sensitivity. To localize nuclear factors associated with starvation sensitivity we generated genotypes that were selectively heterozygous for different combinations of *Zim53;765* autosomes. Twenty *Zim53;765* virgin females were first crossed to 20 males from the double-balancer stock (*wg^Sp-1^/CyO; Dr^1^/TM3, Sb^1^*). Twenty F1 male progeny with both the 2nd chromosome *CyO* and 3rd chromosome *Dr* markers present were sorted over CO_2_ and crossed to 20 *Zim53;765* virgin females in eight replicate 20-female x 20-male crosses. Progeny from this second cross were segregated into four groups based on the presence or absence of the *CyO* and *D*r markers. The four groups were: reconstituted wild-type *Zim53;765*, chromosome 2 heterozygotes (*CyO/765*), chromosome 3 heterozygotes (*Dr/765*) and heterozygotes for both chromosome 2 and 3 (*CyO/765;Dr/765*). Flies from each group were moved to 3 replicate demography cages with approximately 125 flies each. The survival of each group under starvation stress was assayed as described earlier.

### Whole genome sequencing of sensitive and resistant pools from Advanced Intercross Populations

*Population Cages.* Five replicate advanced intercross populations (AIPs) were established between the sensitive (*Zim53;765*) and resistant (*Zim53;315*) lines. Each replicate population was initiated using 25 *Zim53;315* virgin females crossed to 25 *Zim53;765* males, in addition to the reciprocal cross, to yield a total number of 50 male and 50 female parents of each line as the founders of each of the five replicate cages. These parents were introduced into Plexiglas cages (20 x 20 x 20cm) and allowed to lay eggs in four 125mL culture bottles with 30mL of Drosophila medium in each bottle. The four bottles in each cage were left with their tops open for 3 days, and then each bottle was plugged to allow flies to develop. Prior to eclosion, the plugs were removed from each bottle, and all flies were allowed to emerge into the cage and mate at random. Four new culture bottles replaced the old bottles in each replicate cage, and the adults were allowed to lay eggs for 48 hours. After egg laying, adults were cleared from each bottle, plugs were inserted, and cultures were allowed to develop for 7 days following the termination of egg laying, after which the plugs were removed from all bottles to allow eclosion of the next generation into each cage. Population sizes exceeded 1000 individuals after the first generation. All five cages were maintained at large population sizes (>2000 adults) for 10 generations to allow recombination to take place between the sensitive (*Zim53;765*) and resistant (*Zim53;315*) DGRP nuclear genotypes. These two DGRP strains are homozygous for the standard (ST) chromosome arrangement (http://dgrp2.gnets.ncsu.edu/) so this lack of inversions should have allowed free recombination across the genome in these populations.

*Starvation Stress.* The female flies from each generation-10 population cage were collected over CO_2_ and moved to demography cages with a density of approximately 150 flies per cage. The flies were allowed to recover for 7 days with access to high sugar food (Type V: 15% sucrose, 5% Yeast, 2% agar) and then subjected to starvation conditions by replacing the food vials with a 2% agar and water mix. Every six hours from the start of starvation, all the dead flies were collected, flash frozen and stored at −80C. The high sugar treatment enhanced the difference among genotypes in starvation resistance. The genotypic differences in starvation were also evident with moderate or low sugar treatments but mapping experiments were more effective with the high sugar treatment so that was used throughout these analyses.

*Pool-sequencing.* Pools of approximately 50 dead flies were made by combining flies from the two earliest time points when deaths were first observed (24 hour and 30 hour) and the last two time points when the last live flies were present (96 hour and 102 hour). DNA from each replicate pool was isolated using the DNeasy DNA purification kit from Qiagen (Cat No 69506). Each sample was RNase treated prior to DNA elution using a 10-minute RNase A (Cat No 19101) treatment. Libraries were prepared and sequenced using the DNBseq platform by the Beijing Genomics Institute (BGI).

*Estimation of genetic differentiation using Popoolation2.* Raw reads were mapped to the dm6 build of the *Drosophila* genome using *bwa* version 0.7.15 (Li and Durbin 2009). Variants were called using *samtools* version 1.9 (Li *et al*. 2009). Allele frequency differences and estimates of population subdivision (F_ST_) were determined using *Popoolation2* (Kofler *et al*. 2011), and the significance of differences in allele frequencies between the early-dying and late-dying sample across the five replicate populations was determined using the Cochran-Mantel-Haenszel test. P-values were corrected using the Bonferroni method and differences with FDR-corrected p-values < 0.05 were considered to be significant. Linkage disequilibrium was estimated from paired end reads, following the procedure described in (Feder *et al*. 2012). The expectation of this AIP pool-seq experiment is that recombination would segregate major factors responsible for the difference in starvation resistance between these strains so that extreme-phenotype mapping by pooled sequencing might identify chromosomal regions associated with this difference. We acknowledge that 10 generations, moderate populations sizes (∼2000) and founding samples of 50 extreme-phenotype adults may not have the power to generate high resolution recombinants for localizing individual loci, but our intention was to use this as a means to complement other classical mapping approaches (chromosome segregation, marker-recombination assays and deficiency mapping). The use of five replicate populations provided some level of replication that was appropriate for the budget available at the time.

### Chromosome 3 recombination

The chromosome segregation experiments indicated that the majority of the difference in starvation resistance between the lines mapped to chromosome 3 and the pooled sequence analyses of the AIP cages identified two broad candidate regions on chromosome 3, one each on 3L and 3R. We performed recombination mapping using three successive crosses to marked mapping stocks to further localize candidate regions (cross 1, 2, 3; see Fig. S1). For cross 1 we attempted to generate *Zim53;765* chromosome 3 recombinants by crossing 20 *Zim53;765* virgin females to 20 males from a multiply marked recessive stock obtained from the Bloomington Stock Center (BSC_576 = *ru^1^ hry^1^ Diap^th-1^ st^1^ cu^1^ sr^1^ e^s^ ca^1^* hereafter referred to as BSC_576 for convenience). The progeny from this cross however failed to eclose. As an alternative cross 1, we crossed 20 *siI;765* virgin females to 20 *BSC_576* males in an effort to allow recombination across the *DGRP-765* 3^rd^ chromosomes that are present in the *siI;765* stock. The *siI;765* genotype was suspected to be heterozygous for the *DGRP-765* 3^rd^ chromosome and a 3^rd^ chromosome from the *TM3* balancer stock that may have recombined with the *DGRP-765* 3^rd^ chromosome during the construction of the *siI;765* original stock since *siI;765* lacked the *TM3* marker *Sb^1^* (Sb is at 89B4, band 58). This suspicion of balanced heterozygosity was based on the observation that several generations of backcrossing of male DGRP-765 to virgin female *siI;765* did not reduce starvation resistance implying that the *siI;765* carried some heterozygosity and was in some way resistant to the impact of recurrent backcrossing.

For cross 2, 20 F_1_ virgin *siI;765* / BSC_576 females were collected and crossed to 20 males from a multiply marked recessive stock with the same recessive markers as BSC_576 in addition to a dominant marker present on one copy of chromosome 3 balanced over the TM6 balancer (BSC_1711: *ru^1^ hry^1^ Diap^th-1^ st^1^ cu^1^ sr^1^ e^s^ Pri^1^ ca^1^ / TM6B, Bri^1^, Tb^1^*). This cross was done in three replicate 20-female x 20-male vials. F_1_ males from this second cross were collected and segregated into distinct recombinant classes based on the recessive and dominant markers present.

For cross 3, males from each of these recombinant categories were each mated to virgin starvation resistant *siI;765* females and to virgin starvation sensitive *Zim53;765* females to generate offspring with different segments of the 3^rd^ chromosome that was homozygous for the starvation-sensitive *DGRP-765* chromosome. Comparison of the offspring from these two different mitonuclear lines that showed significant differences in starvation (*siI;765* and *Zim53;765*) could further resolve regions of chromosome 3 associated with starvation sensitivity. Due to the presumed heterozygosity of the *siI;765* 3^rd^ chromosome we expected that recombination between the multiple-marked 3^rd^ chromosome with the *siI;765* 3^rd^ chromosomes would be confounded if: 1) the chromosome suspected to be the homologue of the 3^rd^ chromosome in *siI;765* was a broken balancer chromosome, or 2) if that homologue was some other wild type chromosome that had been recombining with the *DGRP-765* 3^rd^ chromosome. This would reduce the efficiency of the recombination mapping as only an unknown fraction of the offspring would have been derived from the actual pairing of the *siI;765* 3^rd^ chromosome derived from DGRP-765 and the recombinant-marked chromosomes derived from the F1 female parent. We acknowledge that this was not ideal but collected the mapping data to see if it provided useful information consistent with the independent chromosome segregation mapping and AIP pool-seq data.

Five classes of recombinant offspring phenotypes were designated based on the visible markers present reflecting different breakpoints along the chromosome: class i*: ru, h*; class ii: *ru, h, Diap, st*; class iii: *ru, h, Diap, st, cu*; class iv: *ru, h, Diap, st, cu, sr*; and class v: *ru, h, Diap, st, cu, sr, e*. Twenty males from each group were crossed to 20 *Zim53;765* virgin females and allowed to mate for 48 hours in three replicate 20-female x 20-male crosses (the *Zim53;765* genotype was confirmed to be homozygous for the DGRP-765 chromosome). The males from each of these crosses were removed and then re-mated to *siI;765* virgin females. Female offspring from each of these two sets of crosses were collected over CO_2_. Sorted by recombinant class and conditioned for starvation assays on high sugar food. Sufficient flies for a starvation stress survival assay were only available from the class i, class ii and class v crosses, so only these were considered. The survival of these females under starvation stress was measured as described above.

### Chromosome 3L Deficiency Screen

The recombination mapping and the AIP pool-seq data implicated chromosome 3L as harboring a major starvation factor, so deficiency mapping was pursued. Twenty males from each deficiency line (Supplemental Table 1) were crossed to 20 *Zim53;765* virgin females in eight replicate 20-female x 20-male crosses. F_1_ females were collected and sorted over CO_2_. Females were separated based on the presence of the respective balancer marker (flies heterozygous for the deficiency and the *765* chromosome do not have the balancer present) and moved to 3 replicate demography cages with approximately 125 flies each, for each genotype. The flies were allowed to recover for 7 days on high sugar food and survival under starvation stress measured as described above.

### iPLA2-VIA Complementation

The chromosome 3L deficiency mapping, in conjunction with the available DGRP sequences, identified *iPLA2-VIA* as a candidate gene based on nucleotide polymorphisms private to *DGRP-765* and starvation sensitivity phenotypes of *iPLA2-VIA* alleles (Lin *et al*. 2018). Deficiencies within that gene were obtained from the Bloomington Stock Center for further complementation mapping: *BSC_80133 = y^1^ w*; iPLA2-VIA^Delta174^* and *BSC_80134 = y^1^ w^1^; iPLA2-VIA^Delta192^* hereafter *80133* and *80134* respectively. These deficiencies were generated by imprecise excision of a P-element from *iPLA-VIA* that removed segments of 5’ noncoding sequence and exons 1-5 and −7 (Lin *et al*. 2018). Twenty *Zim53;765* flies were crossed in reciprocal to 20 flies from each of these two *iPLA-VIA* coding region deficiencies. Each cross was done in replicates of 4. As a control, males from each *iPLA2-VIA* deficiency stock were also crossed to *Zim53;315* females with the expectation that the *DGRP-315* nuclear genome would complement the iPLA2-VIA deficiencies and restore starvation to normal levels. F_1_ females were collected and sorted over CO_2_ and moved to 3 replicate demography cages with approximately 125 flies each. The flies were allowed to recover for 7 days on high sugar food and survival under starvation stress measured as described above.

### Glucose Assay

The free glucose levels, as described in (Tennessen *et al*. 2014), were measured for five genotypes: *Zim53;765*, *Zim53;315*, *iPLA2-VIA^Delta174^*, *Zim53;315* crossed to *iPLA2-VIA^Delta174^* and *Zim53;765* crossed to *iPLA2-VIA^Delta174^*. 120 female flies from each genotype were collected and sorted over CO_2_ and allowed to recover in demography cages affixed with 25×95 mm vials of high sugar food (15% sucrose, 5% Yeast, 2% agar) for five days. Three replicates were done for each genotype. After five days, any dead flies were removed and half of the remaining live flies were removed from the demography cage, immediately flash frozen in liquid N_2_ and stored at −80°C. The food supplied to each demography cage was replaced with 2% Difco Bacto-agar prepared with deionized H_2_0. After 12 hours, dead flies were discarded and the remaining live flies were removed from the demography cage, immediately flash frozen in liquid N_2_ and stored at −80°C. Six frozen flies from each replicate were transferred to a 1.5uL micro-centrifuge tube and homogenized in 120uL of Phosphate Buffered Saline (PBS) using a steel bead and Qiagen TissueLyser II set to 30Hz for 2 minutes. A 10uL aliquot was removed to measure protein content using the Pierce™ BCA Protein Assay Kit (23225). The remaining homogenate was heat treated for 10 minutes at 70°C. The homogenate was diluted 1:4 with PBS. The free glucose was measured with the Glucose Oxidase (GO) kit (MAK097-1KT) by following the protocol for free glucose measurement laid out in (Tennessen *et al*. 2014).

### Data Availability

Drosophila strains, code and raw data are available at the Rand Lab GitHub site for this publication (https://github.com/DavidRandLab/Williams-et-al.-Starvation-Genetics-2021). The authors affirm that all analyzed data necessary for confirming the conclusions of the article are present within the article, figures, tables and GitHub site. Raw sequences for the pool-seq mapping are available at BioProject accession number: PRJNA1130456.

## Results

### Starvation resistance among mitonuclear genotypes

Twelve mitonuclear genotypes from a panel of 72 previously-reported genotypes (Mossman *et al*. 2016) were chosen for starvation assays as described in the Methods section. Each of these 12 mitonuclear introgression lines was subjected to starvation assays on three different isocaloric diets: high sugar food (15% sucrose, 5% Yeast, 2% agar), balanced food (10% sucrose, 10% yeast, 2% agar) or low sugar food (5% sucrose, 15% yeast, 2% agar; referred to as food type V, II and IV, respectively (Zhu *et al*. 2014)). The mitonuclear genotypes on the *DGRP-765* nuclear background were of particular interest because the *DGRP-765* line represents one of the least resistant lines in terms of starvation resistance in the DGRP, and it appeared that introgression of the *siI* mitochondrial DNA haplotype significantly improved resistance to starvation only in the *DGRP-765* nuclear background (Figure 1). To test whether the rescue of starvation resistance mapped to a cytoplasmic or nuclear factor, we performed reciprocal crosses to examine if maternal inheritance of mtDNA was the source of the rescue of starvation sensitivity. When we crossed *Zim53;765* to *siI;765*, the direction of the cross (*Zim53;765 female* crossed *siI;765* males vs. *siI;765* females to *Zim53;765 males*) made no difference to starvation resistance of its progeny. This implied that the *siI* mtDNA haplotype was not the source of the rescue, as would be expected if the trait was being inherited maternally (Fig. S2).

We analyzed the reads from an RNA-sequencing data set from each fly line and compared the genotype calls for each SNP present in *Zim53;765* reads (restricted to the coding regions) and the genotype calls at the same locations in *siI;765*. For the vast majority of these sites, the *siI;765* was heterozygous for the *DGRP-765* allele and an alternative allele (in most cases the reference allele) (Fig. S2A,B). We did a similar comparison of *siI;765* to an alternate mitonuclear genotype (*Zim53;315*) and did not see a similar overlap between the *siI;765* allele and the *Zim53;315* allele (Fig. S2A,B). Based on these results we concluded that the *siI;765* genotypes was heterozygous across the *DGRP-765* chromosomes. This suggested to us that starvation sensitivity in *Zim53;765,* and the two other *mito;765* combinations (*Ore;765* and *siII;765*), were homozygous recessive and the improvement in resistance to starvation seen in *siI;765* was due to its nuclear heterozygosity. Indeed, we observed that the offspring from a *Zim53;765* to *Zim53;315* cross were significantly more resistant to starvation than *Zim53;765* (see Figure 6).

### Chromosome 3 heterozygosity partially suppresses *Zim53;765* starvation sensitivity

Based on the suppression of starvation sensitivity seen in *Zim53;765* heterozygotes, (Zim53;765 x Zim53:315 F1 hybrids) we reasoned that starvation sensitivity phenotypes in *Zim53;765* and other genotypes were homozygous recessive traits and that we could localize the effect locus/loci to chromosomes using chromosome segregation tests and observing which chromosome segregants masked the trait. Virgin females from three different genotypes (the original *DGRP-765, Zim53;765* and *Zim53;315*) were mated to males from the marked double-balancer stock (*wg^Sp-1^/CyO; Dr^1^/TM3, Sb^1^*). F1 males heterozygous for the maternal autosomes paired with the paternal *CyO* and *Dr^1^* markers were mated back to virgin females from the three maternal genotypes, generating the four 2-autosome segregant populations of F1 flies as follows: heterozygous for chromosome 2, heterozygous for chromosome 3, heterozygous for both chromosome 2 and 3, or neither (the latter being reconstituted homozygous wild type 2^nd^ and 3^rd^ chromosomes of the original maternal genotypes: *DGRP-765, Zim53;765* and *Zim53;315*). *DGRP-765* and *Zim53;765* flies heterozygous for chromosome 3 had a ∼200% increase in mean survival relative to the original stock (Figure 2A; Table S2:SummaryTable_figure2a) and flies heterozygous for both chromosomes 2 and 3 realized additional increases in mean survival. Heterozygosity for chromosome 2 alone contributed ∼75% increase (range 50-116%; see Table S2:SummaryTable_figure2a). For Zim53:315, chromosome 3 heterozygosity contributed nothing alone but a small effect in combination with chromosome 2.

**Figure 2.**
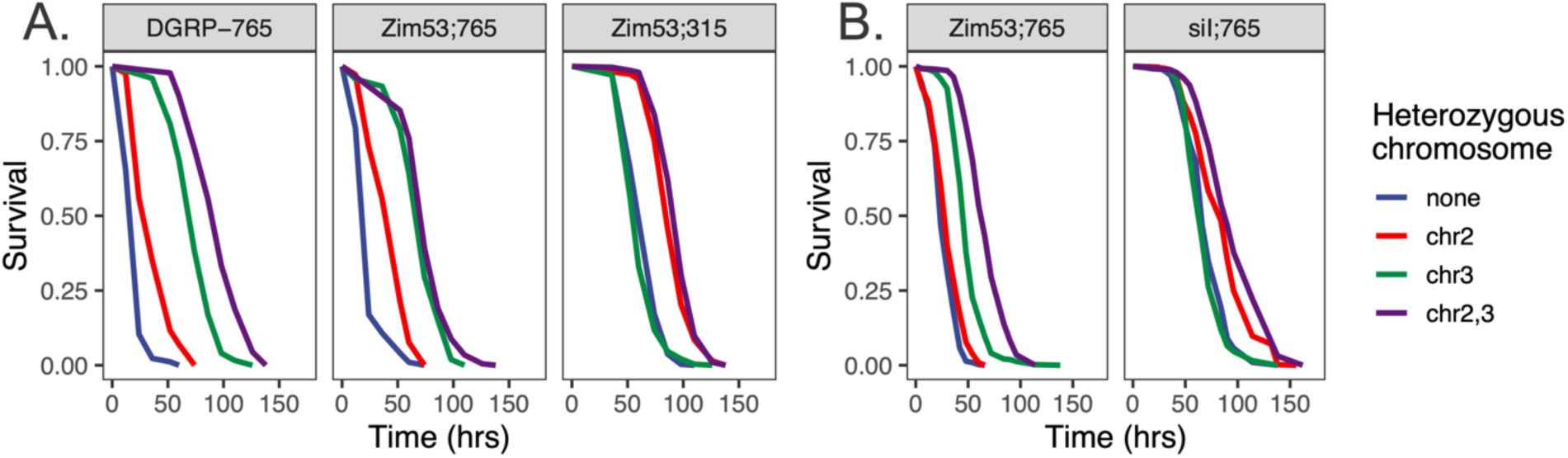
Chromosome 3 suppresses *Zim53;765* starvation sensitivity. (A) *DGRP-765* and *Zim53;765* flies are starvation sensitive while *Zim;315* flies are more resistant. In both *DGRP-765* and *Zim53;765*, heterozygosity for chromosome 3 (chr3) rescues the majority of starvation sensitivity with chromosome 2 heterozygotes (chr2) contributing additional resistance. In *Zim53;315* heterozygosity on chromosome 2 is the primary source of increases in starvation resistance, with little additional contribution from chromosome 3. (B) A replicate experiment with *Zim53;765* shows a similar major effect of chromosome 3 with additional effect of chromosome 2, while the *siI;765* stock with nuclear heterozygosity has increased starvation resistance and distinct chromosomal contributions. Differences in survival between pairs of genotypes were determined using a log-rank test (implemented in the R survival package) and included in Table_S2_Chromosome Segregation Mapping.

A repeat segregation experiment with *Zim53;765* and the heterozygous *siI;765* genotype was conducted and revealed comparable results for *Zim53;765* (80-100% increase from chromosome 3 heterozygosity with additional effects of chromosome 2) and very different chromosomal influences for *siI;765*, as expected from its heterozygous condition (Figure 2B; see Table S2:SummaryTable_figure2b; raw data are in other tabs of Table S2). Taken together, these results suggest that the main effect locus/loci controlling *Zim53;765* and DGRP-765 starvation sensitivity are present on chromosome 3 with a secondary locus/loci present on chromosome 2.

### Sites on chromosome 2R and 3 have significant differences in allele frequency between early and late dying flies

To understand the genetic basis of 765 starvation sensitivity, we sought to identify genetic differences characteristic of starvation sensitive flies. We created replicate advanced intercross populations (AIPs) in which *Zim53;765* was allowed to recombine with *Zim53;315*. We set up five independent populations where *Zim53;765* flies were crossed to *Zim;315* in reciprocal and the populations maintained for 10 generations at population sizes >1000 adults. Following this, females from each population were subjected to starvation stress and dead flies were collected every six hours until all flies were dead. Flies that died during the first three and last two six-hour intervals were separately combined into early- and late-dying pools of adult females (18-24, 24-30 and 30-36 hours vs. 90-96 and 96-102 hours). Total genomic DNA was sequenced from each of the five replicates from these two pooled samples, hereafter ‘early’ and ‘late’ dying pools. Consistent differences in allele frequencies across the early and late dying populations were determined using the Cochran-Mantel-Haenszel test implemented in Popoolation2 (Kofler *et al*. 2011). There was an enrichment in significantly different variants across chromosome 3 and a localized region of chromosome 2R (Figure 3). These results suggest that *Zim53;765* and as a consequence *DGRP-765*, starvation sensitivity is based on variants present on chromosome 3 and chromosome 2. Importantly, these advanced intercross pool-seq analyses are consistent with the chromosome segregation analyses where factors on chromosomes 3 and 2 were responsible for starvation sensitivity of the *Zim;765* genotype.

**Figure 3.**
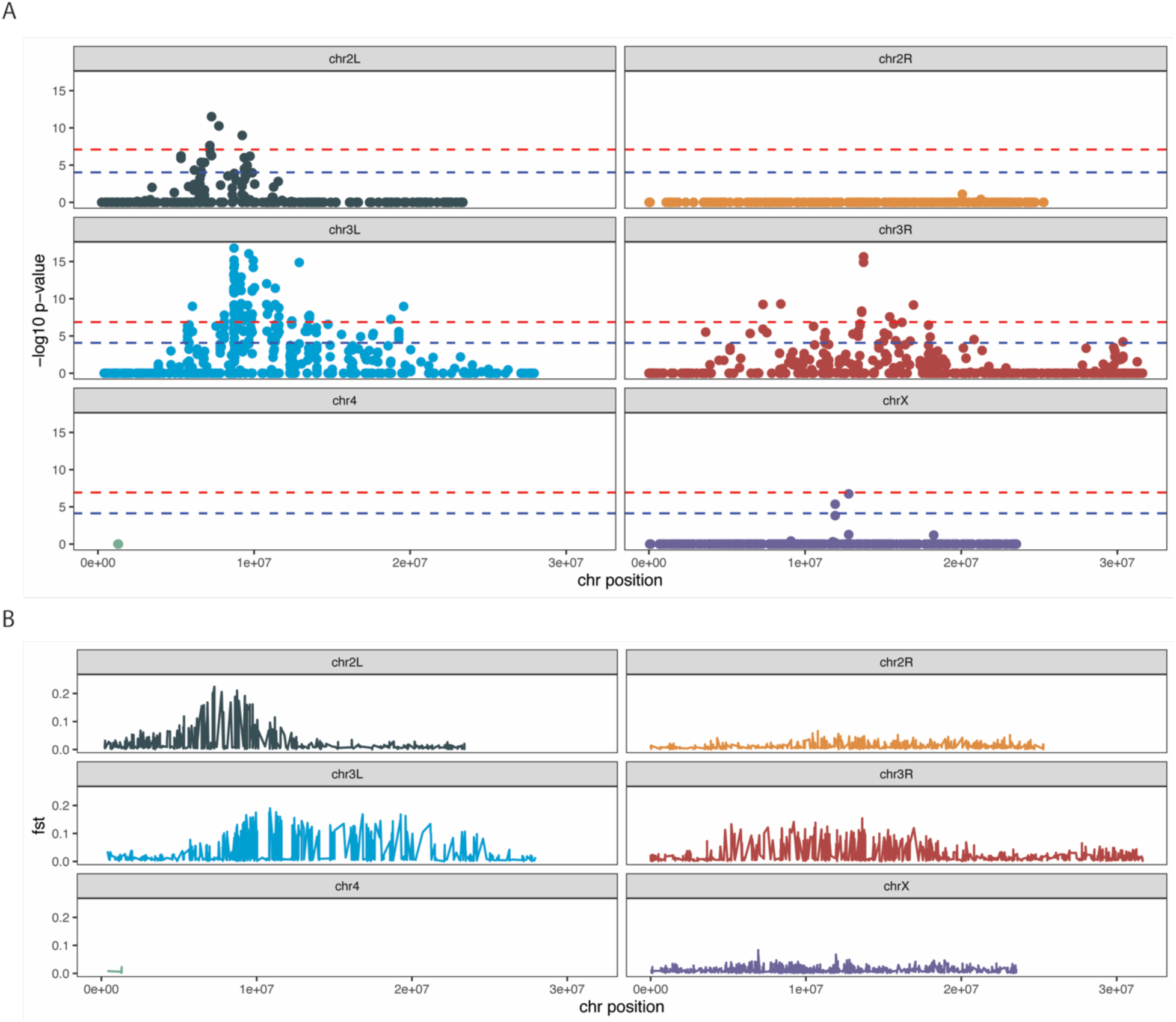
Significant allelic frequency differences between the tails of replicate advanced intercross populations subjected to starvation stress. An advanced intercross population was created by crossing *Zim53;765* and *siI;315* in reciprocal. The hybrids were allowed to recombine freely for 10 generations before being subjected to starvation stress. Dead flies were collected every 6 hours and dead flies from the first 3 timepoints were combined and subjected to sequencing analysis. The same was done with the last two timepoints. In total 5 replicates AIP populations were generated and sequenced. (A) Consistent differences in allele frequencies across replicates were determined using the Cochran-Mantel-Haenszel (CMH) test and Popoolation2. The y-axis is the negative log-transformed, Bonferroni corrected p-value of each SNP (represented by each circle). The x-axis is the position of each SNP along the indicated chromosome. Blue and red lines highlight corrected p-values above 1×10^−5^ and 1×10^−6^ respectively. (B) F_ST_ between pools for each site averaged across replicate populations. The location of elevated FST values is generally concordant with significant CMH test statistics.

### Heterozygosity of chromosome 3L region partially suppresses Zim53;765 starvation sensitivity

To further localize the main effect loci on chromosome 3 we used recombination and deficiency mapping across chromosome 3. As there are over 200 deficiencies spanning chromosome 3, we first generated a series of recombinants with break points at different locations along chromosome 3. Cross 1 used *siI;765* virgin females mated to a multiply marked recessive stock (*ru^1^ hry^1^ Diap^th-1^ st^1^ cu^1^ sr^1^ e^s^ ca^1^* hereafter referred to as BSC_576 for convenience). Cross 2 used heterozygous F_1_ virgin females from cross 1 mated to males from a similarly marked recessive stock with one dominant marker added to distinguish progeny (BSC_1711: *ru^1^ hry^1^ Diap^th-1^ st^1^ cu^1^ sr^1^ e^s^ Pri^1^ ca^1^ / TM6B, Bri^1^, Tb^1^*). The F1 offspring from cross 2 revealed recombinant males that were pooled into three classes of recombinants (classes i, ii, v) that were selected based on the presence of visible recessive markers and sufficient flies for starvation analyses (Figure 4A). Recombinant class i is likely heterozygous for the region of 3L from the telomere up to at least the regions marked by *h,* with recombination breakpoints existing anywhere between h and the *cu* gene. Recombinant class ii is heterozygous for most of 3L and up to *cu* gene on 3R, with breakpoints existing anywhere between the *cu* gene and the *ca* gene. Recombinant class v is likely heterozygous for most of chromosome 3 with breakpoints existing from the *ca* gene up to the telomere of 3R. Other classes of recombinants did not emerge in sufficient numbers to conduct the final mapping crosses. It is important to note that because these recombinants were generated from *siI;765*, the pool of recombinants will carry both the *DGRP-765* alleles from that chromosome which was substituted onto the *D. simulans siI* mtDNA background during original stock construction and alternate, unknown alleles from a chromosome that was likely introduced during the balancer substitution process during original line construction. A *TM3 Sb^1^* 3^rd^ chromosome balancer was used in these crosses and may have recombined with segments of the *DGRP-765* 3^rd^ chromosome during the replacement crosses (see Methods).

**Figure 4.**
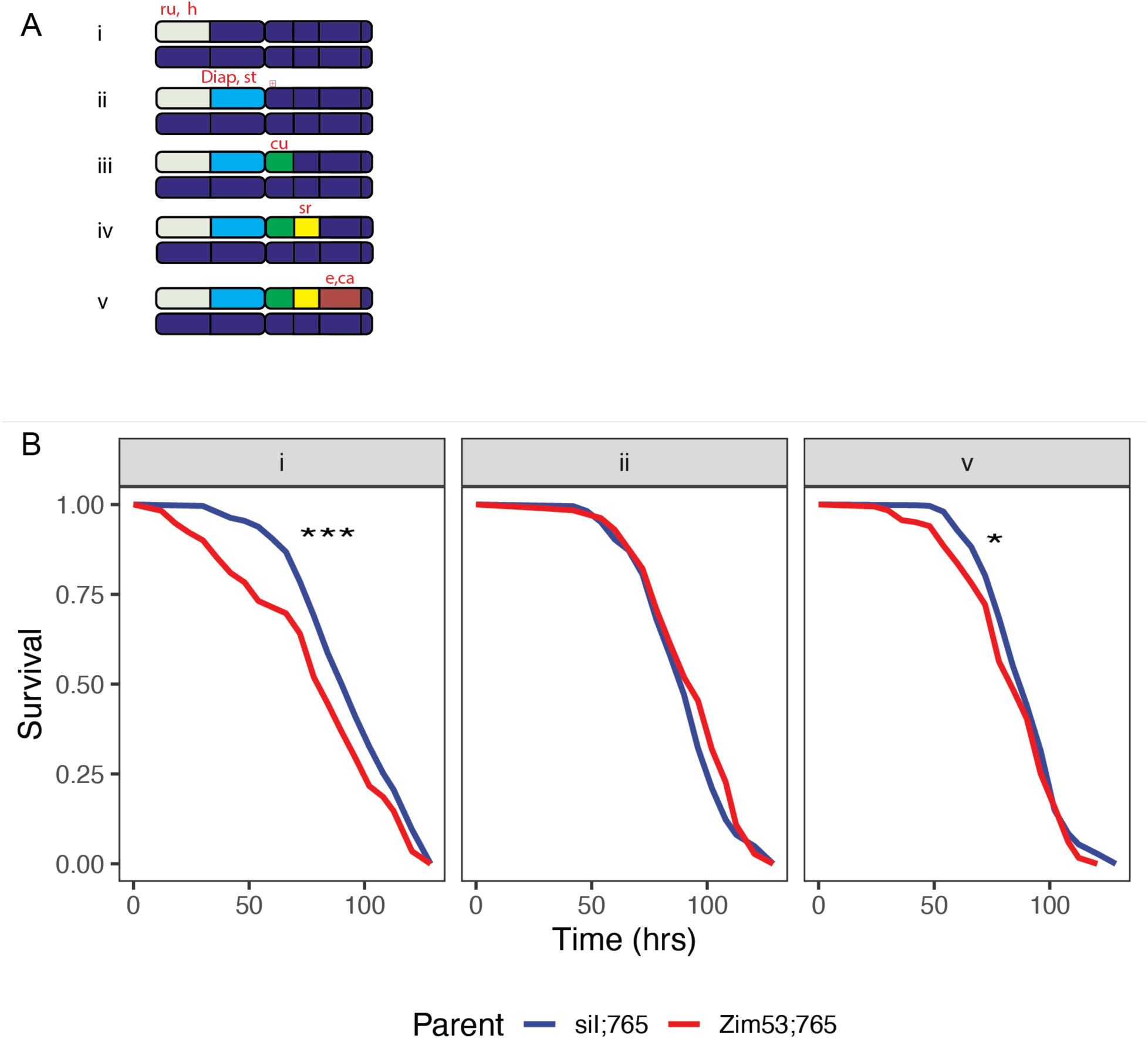
Recombination mapping localizes the majority of *Zim53;765* starvation sensitivity to chromosome 3L. (A) Cartoon representation of recombination breakpoints for each recombinant class generated through meiotic recombination of the chromosome 3 mapping line. (B) Survival of female offspring produced by crossing each recombinant class to *Zim53;765* homozygotes or heterozygotes. Differences in survival were determined using a log-rank test (implemented in the R survival package; *** = p, 0.001, * = p<0.05) and included in Table_S3_Meiotic Mapping Survival.

To reveal the localization, cross 3 used males from the three recombinant classes emerging from cross 2, each of which were mated to virgin *Zim53;765* females, and then the same males were mated to virgin *siI;765* as a control. The starvation sensitivity of the female offspring of the *Zim53;765* and recombinant males was compared to the female offspring of the *siI;765* and recombinant male controls. While we cannot discern genotype at the individual fly level, *DGRP-765* chromosome 3 alleles will be enriched in the progeny from the *Zim53;765* vs recombinant cross, while the alternative will be true for the *siI;765* vs recombinant cross. We expected that flies homozygous for the causal *Zim53;765* allele would die earlier during starvation stress, thus revealing the region of *Zim53;765* chromosome 3 responsible for its starvation sensitivity. Although the heterogeneity of respective offspring populations will mask this effect, the enrichment of the *DGRP-765* allele in the *Zim53;765* vs recombinant cross population and the alternate allele in the *siI;765* vs recombinant cross population should result in appreciable differences in deaths early on during starvation stress. Indeed, we observed a significant difference between the starvation sensitivity of recombinant class i offspring, but not in the case of the class ii offspring; class v offspring showed a small but significant difference (see Figure 4B; Table S3). These results indicate that heterozygosity of a region between the *h* marker of recombinant class i and the *cu* marker of recombinant class ii carries a factor required to suppress *Zim53;765* starvation sensitivity.

### Hemizygosity of 3L: 9812381 – 9899255 prevents partial suppression of *Zim53;765* starvation sensitivity

We employed a deficiency screen across the region between recombinant class i and recombinant class ii markers. The objective of this screen was to identify the region(s) of chromosome 3L that failed to complement *Zim53;765* starvation sensitivity. *Zim53;765* virgin females were crossed to a series of flies deficient for various parts of chromosome 3L (Table S1). The deficient region of each line is maintained over a balancer chromosome so the resulting offspring can be positive for the balancer, indicating complete heterozygosity of chromosome 3, or offspring lacking the balancer are hemizygous across the deficiency region and heterozygous for the rest of chromosome 3. Significant differences in the starvation resistance of the two classes of offspring indicates that heterozygosity of the deficiency region is required for the suppression of starvation sensitivity. We initially screened 15 deficiency lines and identified nine where the starvation sensitivity was not suppressed (failed to complement) in the partial chromosome 3L hemizygote (Table S4A and Fig.S3). Several of the deficiency stock crosses with the most significant difference between Df and wt chromosome pairings shared a common region of chromosome 3 between 3L:9812381 – 3L:9899255 (Fig. S3 and Figure 5), indicating that heterozygosity in this region is required for the suppression of *Zim53;765* starvation sensitivity. Two deficiencies mapped to similar locations but showed conflicting evidence for suppression of starvation sensitivity (Df282 and Df 283; Fig. S3 and Table S1). The latter contained additional genes (lncRNA:CR45878, CG46387) in the proximal region of the deficiency; the functional consequences of these mapping differences were unclear. Three deficiencies on either side of the putative overlap region also showed significant suppression of sensitivity (BSC113, 4470 and 4488). Despite the variation among the deficiency data, they were consistent with the three broader-scale approaches of chromosome segregation, pool-seq and recombination mapping in identifying candidate genes in this region of chromosome 3L.

**Figure 5.**
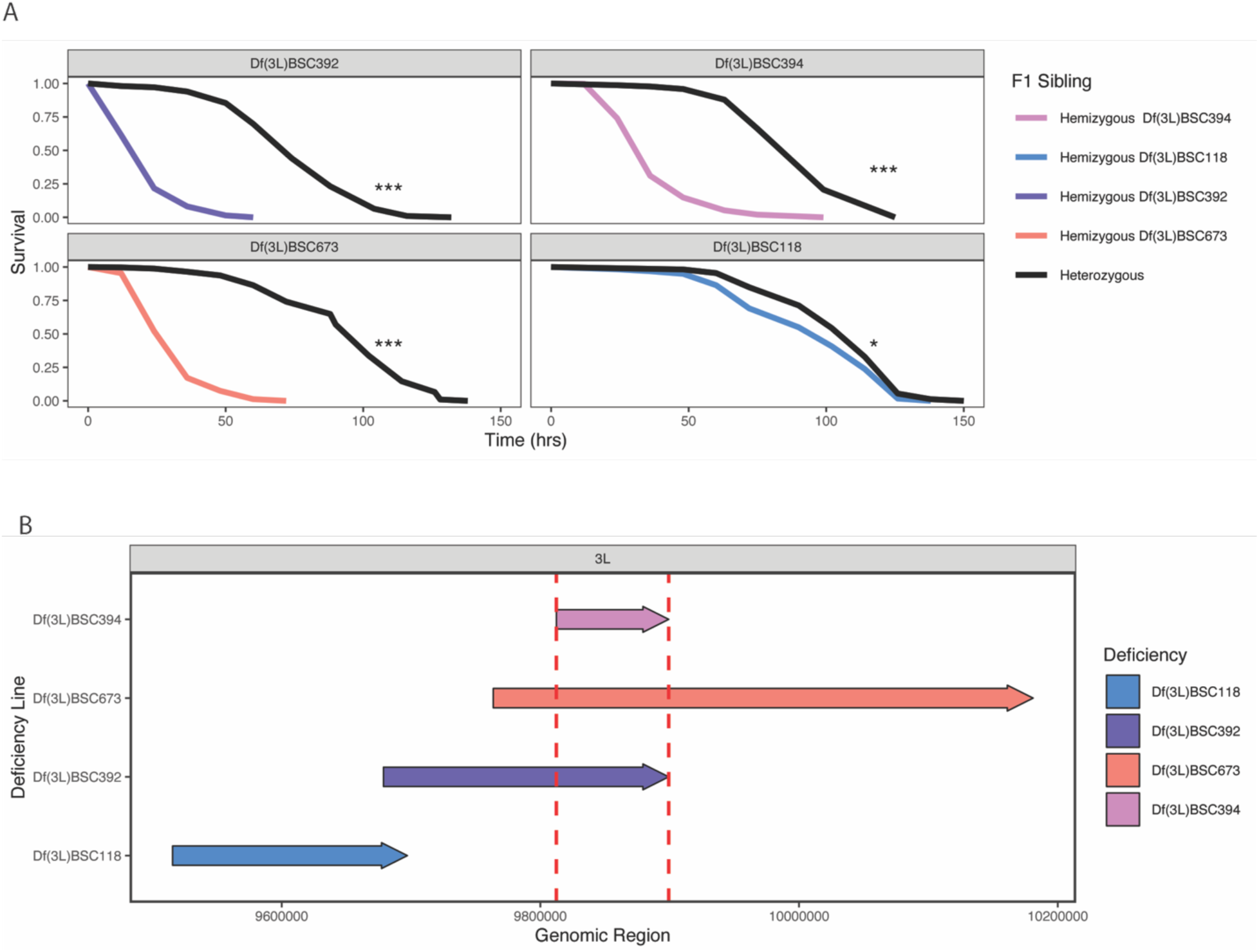
Deficiency mapping identifies the 3L: 9812381 – 9899255 region as a component of *Zim53;765* starvation sensitivity. (A) Survival of offspring from the cross between female *Zim53;765* and male deficiency stocks following starvation stress. Heterozygous refers to progeny that inherited the marked balancer chromosome, hemizygous refers to progeny that inherited the deficient chromosome. Line numbers correspond to Bloomington Stock Center names (see table S1). Differences in survival were determined using a log-rank test (implemented in the R survival package) and included in Table S4_Deficiency mapping: summary and statistics tabs. (B) Linear representation of hemizygous regions of respective deficiency lines. Highlighted region is common to the three deficiency regions which failed to suppress *Zim53;765* starvation sensitivity. See Fig. S3 for additional crosses

### Variation in iPLA2-VIA as candidates for *Zim53;765* starvation sensitivity

Next, we sought to identify *Zim53;765* variants in the chromosome 3L region that might underlie its sensitivity to starvation. We utilized the available genomic sequencing data of each DGRP line to identify nucleotide variants unique to the *DGRP-765* sequence (Mackay *et al*. 2012). As the causal variant is deleterious, we focused our initial analysis on low frequency variants (present in less than 10 % of the 200 DGRP2 sequences). We identified 83 variants which fit this frequency cutoff and only thee SNPs were predicted to lead to non-synonymous changes (Fig. S4 and Table 2). A C-to-A mutation, predicted to result in a G-to-V amino acid residue change in the conserved GXGXXG DNA binding motif of the phospholipase *iPLA2-VIA*, appeared as a strong candidate. Mutations in *iPLA2-VIA* have previously been shown to cause starvation sensitivity and reduced lifespan in *Drosophila (*Kinghorn *et al*. 2015; Kinghorn and Castillo-Quan 2016; Lin *et al*. 2018). In addition, PolyPhen-2 predicted the mutation as probably damaging, with a score of 0.999 out of 1 (Adzhubei *et al*. 2010). Based on this analysis, we performed finer resolution deficiency mapping in the *iPLA2-VIA* gene.

**Table 2.** Sequence details of candidate SNPs in the genomic region identified by classical mapping approaches. See text and Fig. S4 for details.

| Chr. | start | end | REF | ALT | CONSEQUENCE | REFDNA | VARDNA | REFAA | VARAA | SYMBOL |
| --- | --- | --- | --- | --- | --- | --- | --- | --- | --- | --- |
| chr3L | 9824166 | 9824166 | C | T | nonsynonymous | CCA | TCA | P | S | CG43897 |
| chr3L | 9833935 | 9833935 | G | A | synonymous | AGC | AGT | S | S | RasGAP1 |
| chr3L | 9854881 | 9854881 | C | A | nonsynonymous | GGC | GTC | G | V | iPLA2-VIA |
| chr3L | 9880027 | 9880027 | A | G | synonymous | TTT | TTC | F | F | Taf2 |
| chr3L | 9880564 | 9880564 | G | T | synonymous | ATC | ATA | I | I | Taf2 |
| chr3L | 9881643 | 9881643 | C | A | nonsynonymous | GAG | GAT | E | D | Taf2 |
| chr3L | 9884329 | 9884329 | A | T | synonymous | ATA | ATT | I | I | CalpB |
| chr3L | 9884605 | 9884605 | A | G | synonymous | GGA | GGG | G | G | CalpB |

### *iPLA2-VIA^Del174^* fails to complement *Zim53;765*/ DGRP-765 starvation sensitivity and glucose homeostasis

To determine if the starvation phenotype of *Zim53;765* and as a consequence *DGRP-765*, results from a defective copy of *iPLA2-VIA*, we sought to determine if the *Zim53;765* copy of *iPLA2-VIA* gene could complement *iPLA2-VIA* null mutations. We crossed *Zim53;765* virgin females to two previously characterized *iPLA2-VIA* null lines (*iPLA2-VIA^Del174^*, *iPLA2-VIA^Del192^),* generated by imprecise excision of a mobile element; Table S1 (Lin *et al*. 2018). To rule out the potential effects of mtDNA haplotype on starvation resistance, we also utilized the original *DGRP-765* strain obtained from the Bloomington Stock center. We found that progeny from *DGRP-765* crossed to *iPLA2-VIA^Del174^* or *iPLA2-VIA^Del192^*were also starvation sensitive (Figure 6). In contrast, crossing *iPLA2-VIA^Del174^* or *iPLA2-VIA^Del192^* to the starvation resistant strain *Zim53;315* resulted in the suppression of starvation sensitivity (i.e., restored starvation resistance; Figure 6).

**Figure 6.**
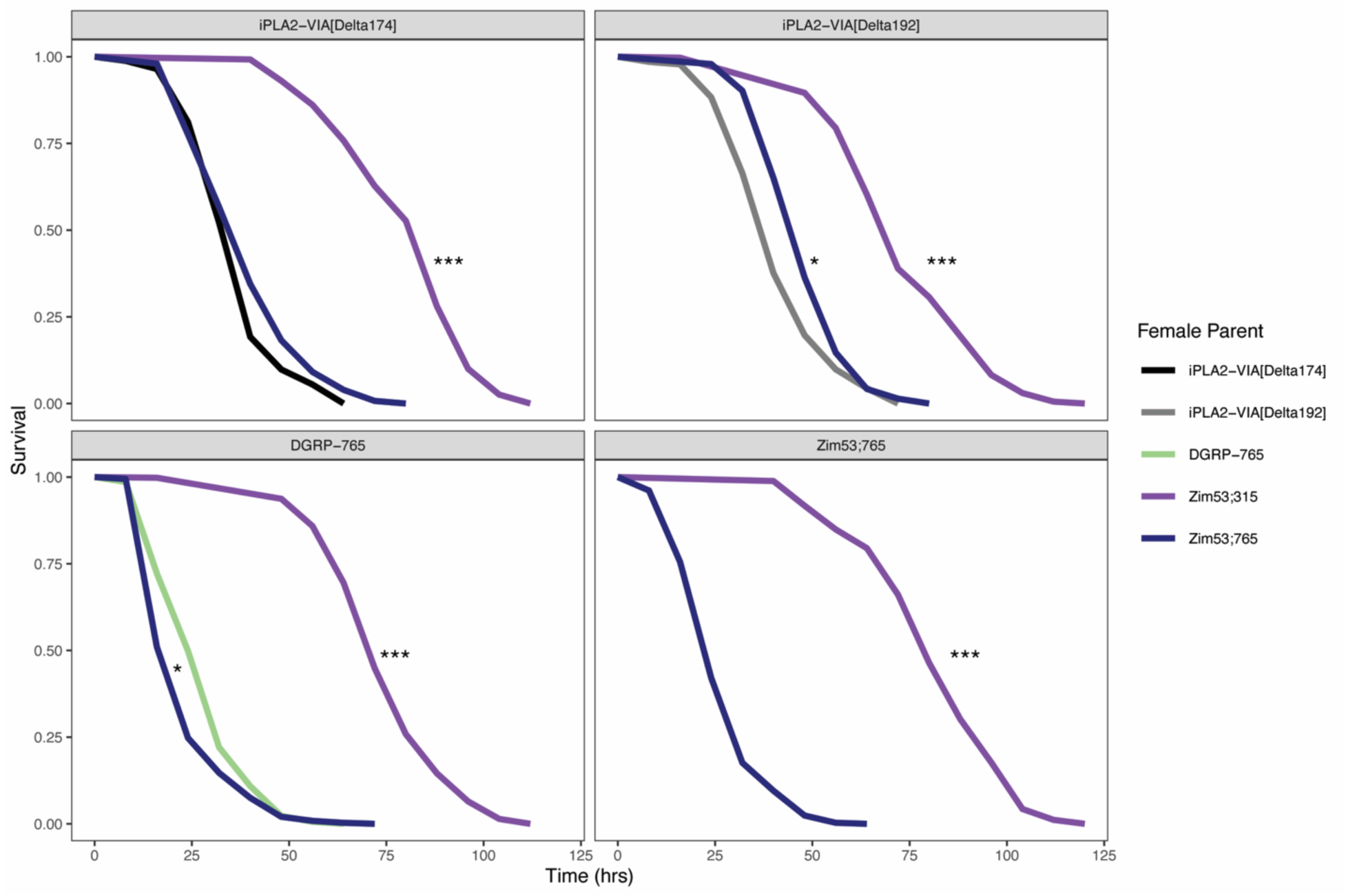
*Zim53;765* fails to complement iPLA2-VIA null lines. Survival of female offspring from indicated crosses following starvation stress. Each panel represents the male parent, and colored lines identify the female parent. *Zim;765* is starvation sensitive as a homozygote and as a heterozygote over two different *iPLA2-VIA* deficiency null alleles. Differences in survival were determined using a log-rank test (implemented in the R survival package) and included in Table S5_iPLA-VIA complementation).

### Depletion of glucose and glycogen are associated with starvation sensitivity in *Zim53;765* and *DGRP-765*

Ablation or pharmaceutical inhibition of calcium independent phospholipases have been shown to reduce insulin secretion in response to a glucose challenge (Song *et al*. 2005; Ali *et al*. 2013). This raised the question of whether *Zim53;765* and *iPLA2-VIA* share defects in glucose metabolism. We observed that following 12 hours of starvation, *Zim53;765* and the original *DGRP-765* line significantly deplete their levels of free glucose and glycogen in contrast to *Zim53;315,* suggesting improper regulation of glucose during acute starvation (Figure 7). Interestingly, this phenotype was not observed in *iPLA2-VIA^Del174^* homozygotes (Figure 7). However, we found that progeny from *Zim53;765* crossed to *iPLA2-VIA^Del174^*also significantly deplete their available free glucose following 12 hours of starvation (Figure 7; Table S6a,b,c). Taken together, these results indicate shared genetic bases of *iPLA2-VIA* and *Zim53;765* in starvation sensitivity but distinct roles in glucose metabolic effects on starvation. Notably these effects follow a week of conditioning on high sucrose foods that may influence the time course of sugar depletion during starvation and its impact on survival during starvation.

**Figure 7.**
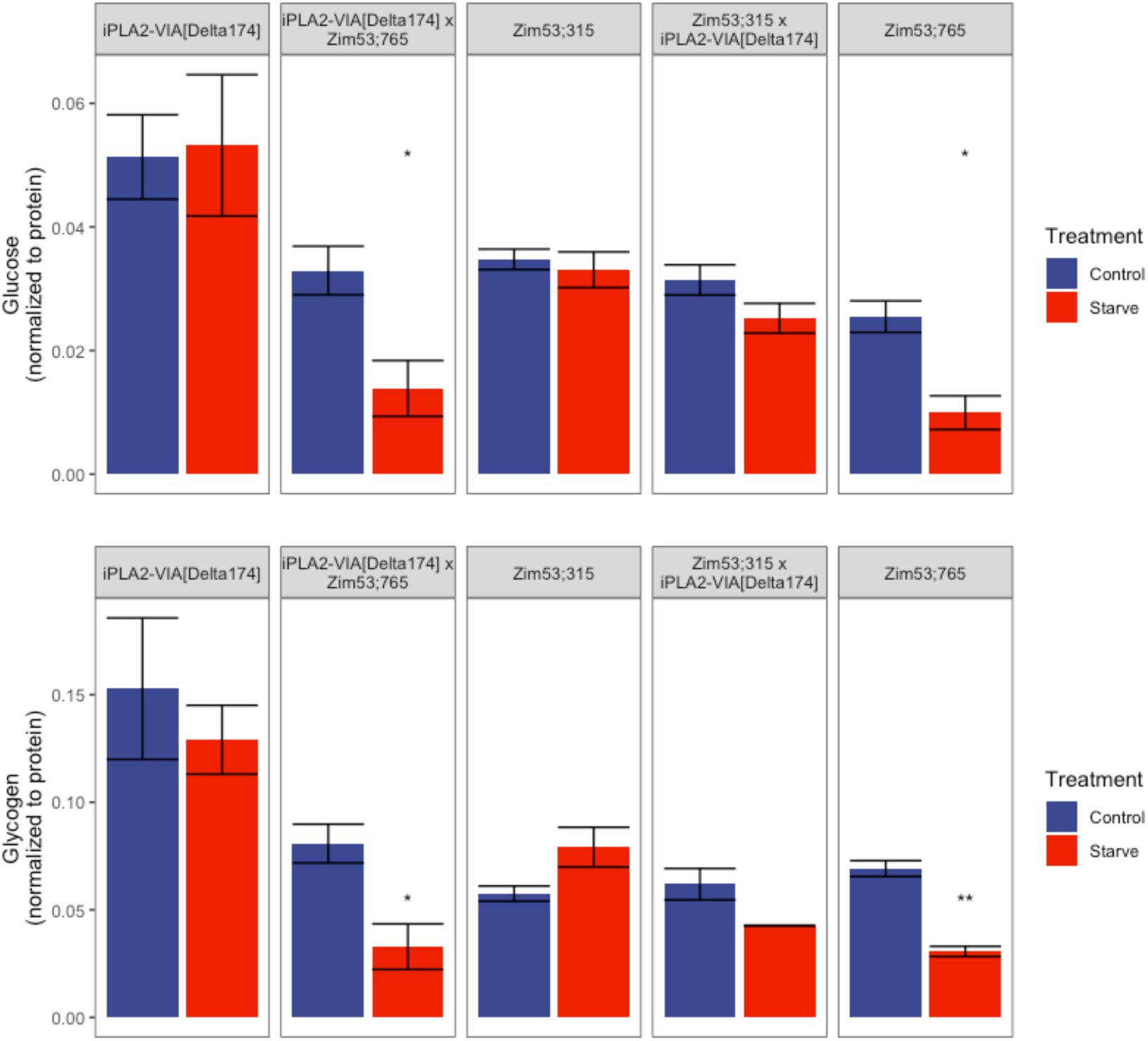
iPLA2-VIA[Delta174] fails to complement glucose and glycogen depletion phenotype of *Zim53;765*. The free glucose and total glycogen levels were measured for *Zim53;765*, *Zim53;315*, iPLA2-VIA[Delta174], *Zim53;315* crossed to iPLA2-VIA[Delta174] and *Zim53;765* crossed to iPLA2-VIA[Delta174]. Values in the un-starved (control) was compared to the values in the starved samples (12 hours) for 3 replicates of 6 flies using a student’s t-test. Significant differences are indicated by * <0.05, ** <0.01.

## Discussion

In the current study we sought to identify the genetic basis of a striking difference in starvation resistance between one particular genotype among a set of related mitonuclear genotypes (see Figure 1). Given the central role that mitochondria play in both anabolic and catabolic processes related to energy stores, the possibility that combinations of mitochondrial and nuclear genes might contributed to quantitative variation for starvation resistance warranted additional genetic analysis. This is further motivated by the highly polygenic nature of starvation resistance and its correlation with other fitness and metabolic traits (Service and Rose 1985; Chippindale *et al*. 1996; Harshman *et al*. 1999; Hardy *et al*. 2018; Everman *et al*. 2019). Our early crosses ruled out an effect of the mtDNA on the strong differences in starvation between *siI;765* and *Zim53;765* which refocused the analyses to the genetic architecture of starvation resistance between distinct nuclear genomes. Because the nuclear genomes used to build the mitonuclear lines were part ofthe well-characterized DGRP we sought to use the classical tools of the Drosophila model and the genomic information available for the DGRP to carry out a quantitative trait locus search between two distinct homozygous lines from the DGRP. The motivation was twofold: the difference between the sensitive mitonuclear lines (in *DGRP-765* backgrounds) and resistance lines (in *DGRP-315* or -*820* backgrounds) was very distinct, and the sensitive *mitonuclear-765* lines also appeared less sensitive to the high sugar:protein ratio in the food used prior to the starvation assay (Figure 1). Applying forward genetic approaches to this classic quantitative genetic system offered the opportunity to test the hypothesis that starvation sensitivity was associated with sugar metabolism.

### Polygenic basis of *Zim53;765* starvation sensitivity

The chromosomal segregation analyses and the Advanced Intercross Population (AIP) pool-seq analyses show parallel results at a chromosomal scale and further confirm the polygenic basis of the differences between the *DGRP-315* and *-765* genomes. Figure 2 shows a clear main effect of chromosome 3 and some additive or epistatic effects from chromosome 2. Chromosome 3 heterozygosity was necessary to suppress *Zim53;765* starvation sensitivity but heterozygosity of chromosome 2 and 3 together further improved starvation resistance. Notably, chromosome 2 heterozygosity in isolation had limited effect on suppressing *Zim53;765* starvation sensitivity (Figure 2A, B). This indicates that the chromosome 2 locus/loci acts as a modifier that genetically interacts with the chromosome 3 locus/loci. Importantly, *Zim53;315* has a distinct genetic basis of starvation resistance, with chromosome3 having little effect (Figure 2A)

The pool-seq results from the AIP design (Figure 3) show corresponding patterns for chromosomes 2 and 3: strong evidence on chromosome 3 for allele frequency differences between early-and late dying pools of flies, but limited evidence for this on chromosome 2. Figure 2 shows a weak signal of starvation suppression by chromosome 2 for the very last-dying flies but it is not significant. This could reflect heterozygosity of the few loci that just pass significance on chromosome 2L in the pool-seq data (Figure 3), but this remains speculative.

We note that the AIP analyses involved recombination between the focal line at the extreme end of the starvation resistance spectrum (*Zim53;765*, most sensitive) and a line more central in the distribution of starvation scores (Zim;315, with the DGRP-315 nuclear chromosomes). Though a compelling case could be made for recombining Zim;315 with a line at the opposite end of the distribution (most resistant; e.g., Zim;820 with the *DGRP-820* nuclear chromosomes), our starvation data did not show a large difference between the -*315* and -*820* mitonuclear genotypes, but the -*315* mitonuclear genotypes had somewhat tighter variance of starvation scores on the different sugar:protein diets, so we chose that background (Figure 1). The AIP pool-seq data show strong allelic differentiation across one broad peak on chromosome 3L and additional peaks on 3R, with fewer significant peaks on chromosome 2L (4 SNPs above threshold). The resolution of these analyses is admittedly low and resolving specific candidate loci was not an expected goal of the AIP pool-seq. It is likely that many of these significant variants are in linkage disequilibrium since the populations recombined over a period of only 10 generations. Nevertheless, the clustering of significant variants on chromosomes 2 and 3 are consistent with a polygenic basis for *Zim53;765* starvation sensitivity. The AIP analyses served the important goal of confirming the results of the chromosome segregation analyses and identifying where to focus subsequent higher resolution recombination and deficiency mapping.

### Dissecting the 3L candidate region

We sought to narrow the target region, by generating recombinants across chromomere 3 between *Zim53;765* and the multiply marked recessive stocks, as this would provide a more localized region for deficiency analyses. We however failed to do this directly as crossing *Zim53;765* females to males from each of the multiple marked recessive stocks failed to yield an appreciable number of offspring. We use the *siI;765* to generate recombinants because *siI;765* was constructed using balancer substitutions and backcrosses to *DGRP-765* and thus likely carried *DGRP-765* chromosomes, and it produced sufficient offspring to proceed with the remaining two recombination crosses. Despite learning that *siI;765* was a hybridized version of *765* nuclear chromosomes, a fact confirmed by comparing the alleles of *Zim53;765* variants to *siI;765*’s, it remained possible that the recombinant offspring would help localize genetic effects between chromosome arms 3L or 3R, which proved true. The 15 different deficiencies used for mapping were chosen to span the regions of chromosome 3L that were most likely to carry a candidate gene or genes contributing to the starvation differences between the DGRP chromosomes present in *Zim;765* and *Zim;315*. The preponderance of the evidence from these data supported the hypothesis that heterozygosity of the region of 3L 9812381 – 9899255 was required for suppression of *Zim53;765* starvation sensitivity, with deficiencies BSC392, -394 and -673 being clear examples (Figures 5 and S3). Again, however, the deficiency data included some inconsistent results. When *Zim53;765* was crossed with Df(3L)BSC282, there was only a modest difference in the starvation resistance of the heterozygous and homozygous progeny, and when crossed with Df(3L)BSC283, there was no statistical difference in the starvation resistance (Fig. S3). The two additional genes missing in Df(3L)BSC283 relative to Df(3L)BSC282 may explain these conflicting data, but neither gene has any known function reported in Flybase (lncRNA:CR45878, CG46387).

### Complementation points to Zim53;765 iPLA2-VIA

There are over 30 genes in the candidate regions defined by 3L 9812381 – 9899255 (see Table S1). Of all the variants in this region, the nonsynonymous mutation present in the calcium-independent phospholipase iPLA2-VIA immediately stood out, due to the location of the mutation in the functional domain of the enzyme, and the phenotypes previously associated with iPLA2-VIA loss of function. Mutations in the human homolog PLA2G6 gene are characteristic of multiple autosomal recessive disease (Guo *et al*. 2018). The amino acid mutation in the Drosophila iPLA2-VIA protein that is private to *DGRP-765* lies in the GXGXXG consensus nucleotide binding motif similar to those found in other protein kinases. This motif contributes to the binding of ATP to iPLA2-VIA which is essential for its catalytic function (Ramanadham *et al*. 2015). Several studies have independently characterized starvation sensitivity in *iPLA2-VIA* mutants (Kinghorn *et al*. 2015; Iliadi *et al*. 2018; Lin *et al*. 2018; Mori *et al*. 2019). In each case, targeted mutations were introduced in *iPLA2-VIA* to generate null models, with the goal of better understanding its role in neurodegeneration. These studies identified aberrant lipid metabolism, lipid peroxidation and mitochondrial dysfunction as potential causes of stress sensitivity and neurodegeneration. We should not necessarily expect point mutations in *iPLA2-VIA* to universally have the same effect as these deletion loss of function models, as mutations in PLA2G6 have varying effects on catalytic activity (Engel *et al*. 2010). However, we can expect that if mutation(s) in *iPLA2-VIA* do in fact underlie *Zim53;765* starvation resistance, an *iPLA2-VIA* null should fail to complement *Zim53;765* starvation sensitivity. We showed that two separate nulls generated by (Lin *et al*. 2018) do in fact fail to complement *Zim53;765* starvation sensitivity, indicating that natural variants in iPLA2-VIA contribute some of the starvation sensitivity of *Zim53;765* and *DGRP-765*.

The correlation between starvation resistance and lipid content has been demonstrated across species and as associated responses to selection for starvation resistance (Chippindale *et al*. 1996; Harshman *et al*. 1999; Hardy *et al*. 2018; Everman *et al*. 2019). While our current study does not have details on lipid content, there may be a link to the glucose and glycogen utilization we report (Figure 7), and to other possible candidate genes in the 3L 9812381 – 9899255 region: there are several insulin-like peptides (*Ilps*) ∼40-100kb distal to the 3L 9812381 – 9899255 that show polymorphism among the DGRP lines. Some phospholipases have been associated with modulation of insulin secretion in response to glucose by releasing arachidonic acid (AA), and exogenous AA supplementation is sufficient to promote insulin release from rat islet cells (Metz 1988a). This insulin secretion is believed to promote phospholipid hydrolysis and accumulation of phospholipid-derived intermediates (Metz 1988b). This turnover of phospholipids could promote insulin release, creating an interesting dynamic between glucose and lipid metabolism, with phospholipases at the center. The premature depletion of glucose and glycogen in *Zim53;765* in the early stages of starvation could be one consequence of the SNP(s) in iPLA2-VIA or an interaction with *Ilps*. Consistent with this, we found that *iPLA2-VIA^Del174^* fails to complement the premature glucose and glycogen depletion phenotypes associated with *Zim53;765* (Figure 7). What is especially interesting is that the iPLA2-VIA null itself does not display this phenotype. This raises the possibility that affected pathways which ultimately result in starvation sensitivity differ between the deletion mutation of *iPLA2-VIA^Del174^* and the point mutation unique to *Zim53;765*. It will be an interesting target of a future study to characterize the additional molecular bases underlying the starvation sensitivity of *Zim53;765,* as the amino acid mutation in the conserved GXGXXG motif of *iPLA2-VIA* could potentially highlight novel functions of *iPLA2-VIA*.

## Acknowledgments

The authors thank the HHMI Summer Fellows Julia Dewey, Tyler Devlin, Brian Franklin, Cynthia Hale-Phillips, Matthew P. McAteer, Zemplen Pataki, Chén Yé, and Denise Yoon who conducted the initial screen for mito-nuclear interactions affecting starvation resistance, which provided the impetus for this work. Dr. Jim Mossman assisted with the analyses of DGRP polymorphisms. Yevgeniy Raynes and Leah Darwin provided helpful edits for R code.

---

**Supplemental Figure S1.**
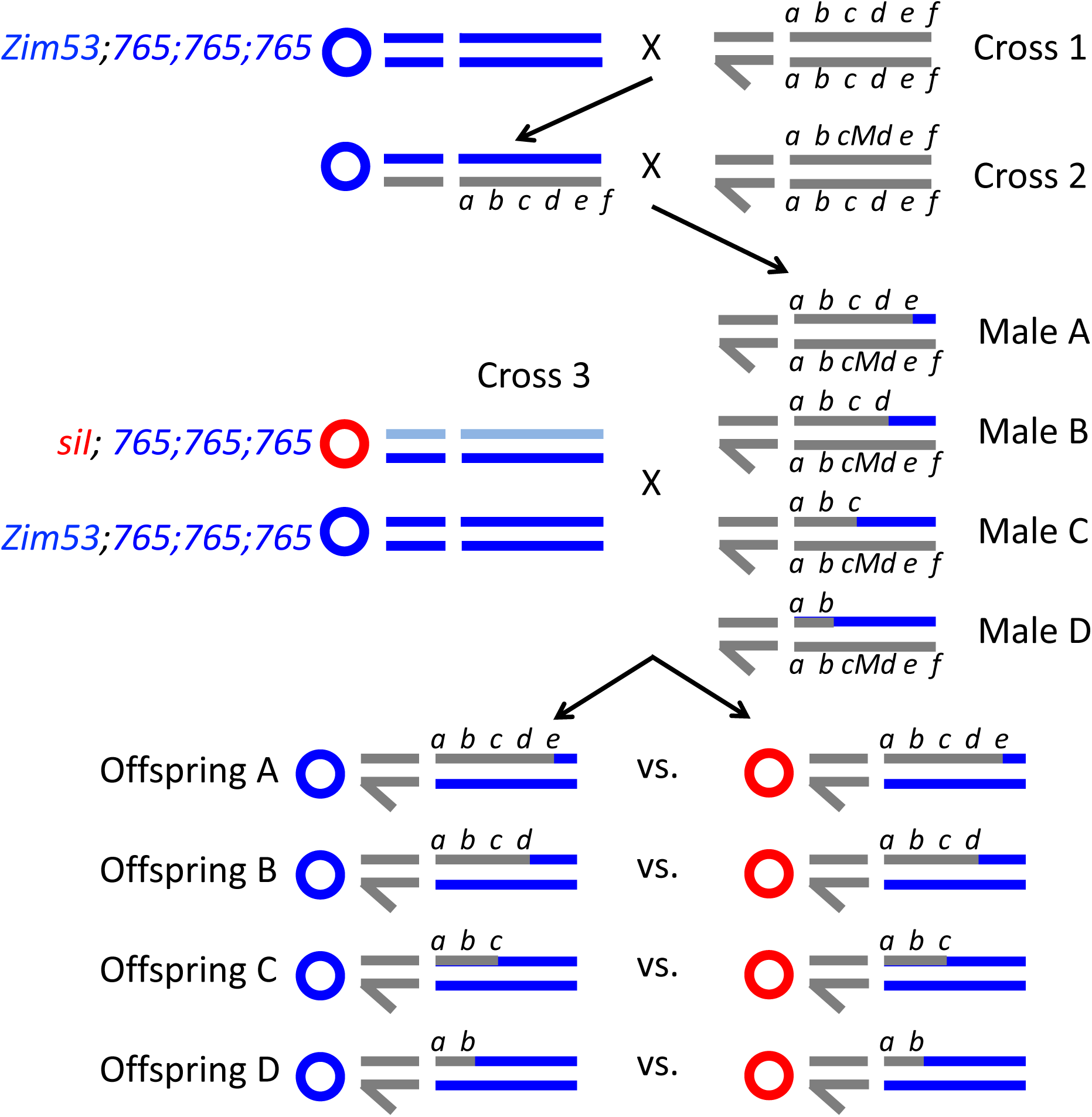
Recombination scheme for meiotic mapping. The scheme includes three sequential crosses using the focal lines described in the text and two multiply marked stocks to resolve recombination breakpoints. Note that *Zim53;765* is homozygous for DGRP-765 chromosomes and is starvation sensitive while *siI;765* is starvation resistant but has nuclear heterozygosity is starvation resistant (see Figure 1).

**Supplemental Figure S2.**
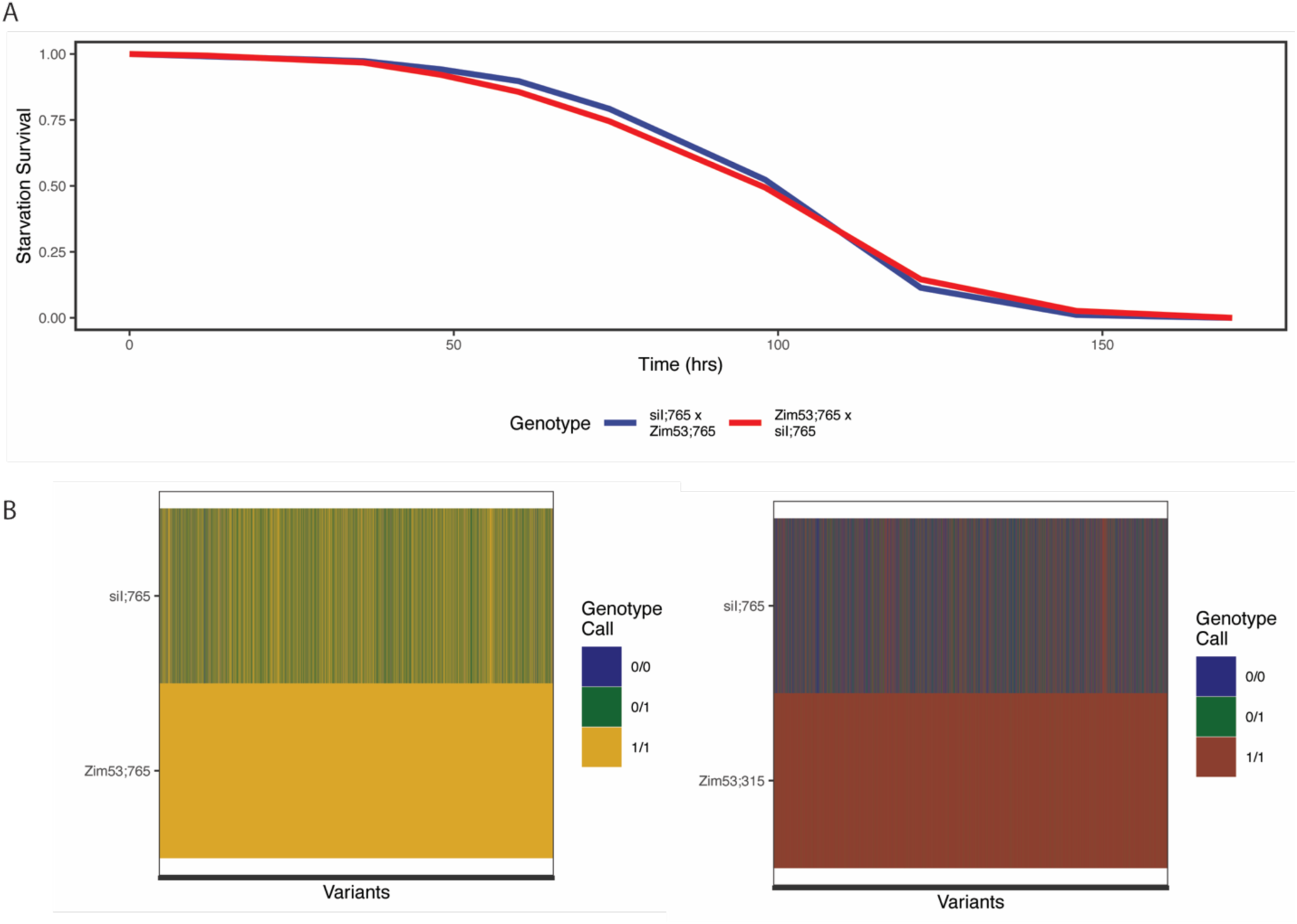
*siI;765* is a hybrid of DGRP-765. (A) Survival of female offspring of *Zim53;765,* and *siI;765* crossed in reciprocal. There is no statistical difference in survival based on a log-rank test for significance. (B and C) Genotype of *siI;765* at homozygous variant sites in RNA sequencing reads from (C) *Zim53;765* and (D) *Zim53;315* Green indicates that *siI;765* was heterozygous for both the variant and the reference allele, blue indicates that *siI;765* was homozygous for the reference allele.

**Supplemental Figure S3.**
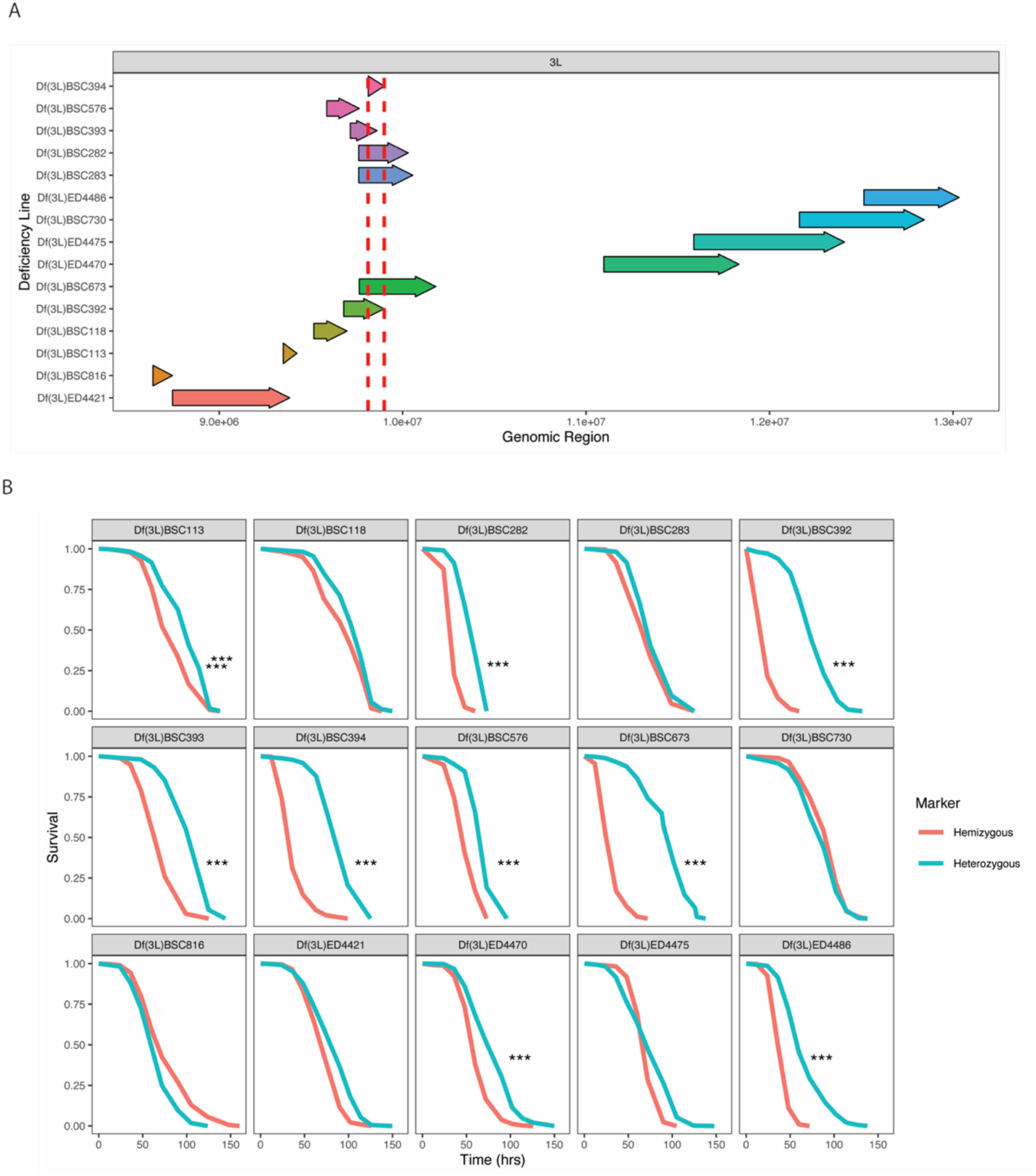
(A) Linear representation of deficient region of respective deficient lines in region identified in meiotic recombination mapping. (B) Survival of offspring from the cross between *Zim53;765* and the respective deficient male following starvation stress. Differences in survival were determined using a log-rank test (implemented in the R survival package) and included in TableS4_Deficiency mapping. Heterozygous refers to progeny that inherited the balancer chromosome, hemizygous refers to progeny that inherited the deficient chromosome. Line numbers correspond to Bloomington Stock Center names.

**Supplemental Figure S4.**
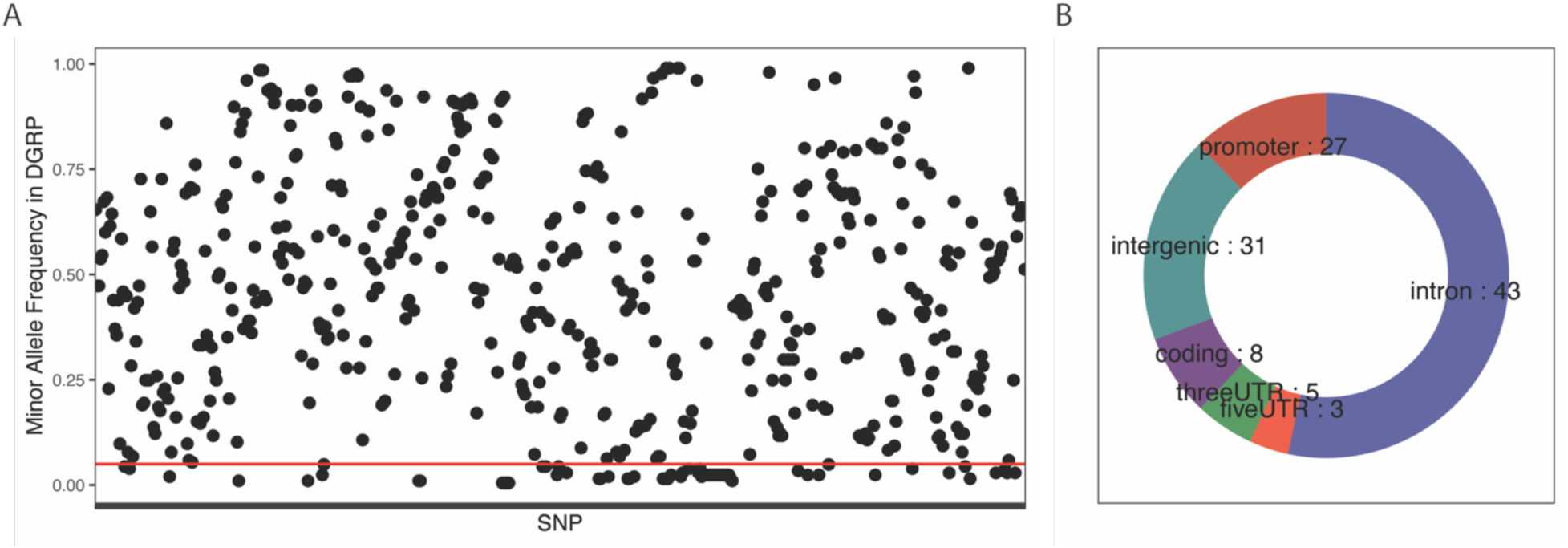
Frequency of DGRP-765 variants among the Drosophila Genetic Reference Panel. (A) Allele frequency of *DGRP-765* variants in the 3L candidate region among the lines of the Drosophila Genetic Reference Panel. Variants found below the red line are present in less than ten percent of the lines. (B) Proportions of low frequency (<10%) variants in different functional categories for in the 3L candidate region present in DGRP-765.

